# Prebiotically Plausible Activation Chemistry Compatible with Nonenzymatic RNA Copying

**DOI:** 10.1101/2020.05.13.094623

**Authors:** Stephanie J. Zhang, Daniel Duzdevich, Jack W. Szostak

## Abstract

The nonenzymatic replication of ribonucleic acid (RNA) oligonucleotides may have enabled the propagation of genetic information during the origin of life. RNA copying can be initiated in the laboratory with chemically activated nucleotides, but continued copying requires a source of chemical energy for *in situ* nucleotide activation. Recent work has illuminated a potentially prebiotic cyanosulfidic chemistry that activates nucleotides, but its application to nonenzymatic RNA copying remains a challenge. Here we report a novel pathway that enables the activation of RNA nucleotides in a manner that is compatible with template-directed nonenzymatic polymerization. We show that this pathway selectively yields the reactive imidazolium-bridged dinucleotide intermediate required for nonenzymatic template-directed RNA copying. Our results will enable more realistic prebiotic chemical simulations of RNA copying based on continuous in situ nucleotide activation.

---

RNA is a leading candidate for the primordial genetic polymer because of its potential to act as both a hereditary and functional biomolecule ^1–3^. The emergence of life in the RNA World would have required nonenzymatic RNA replication to propagate genetic information prior to the emergence of more elaborate ribozyme-catalyzed replication machinery ^4–6^. Primer extension is a model of RNA copying in which nucleotides **1** are added to a primer when guided by a template sequence (Fig. 1) ^7–9^. Nonenzymatic primer extension relies on activation of the mononucleotide phosphate groups ^10–14^.

**Figure 1.**
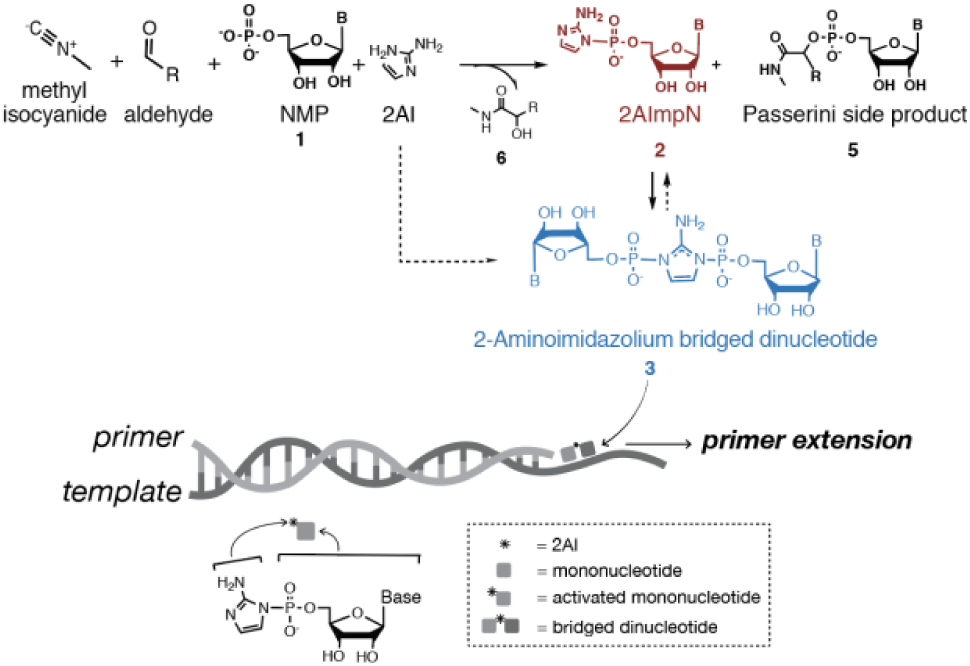
The components of nonenzymatic RNA primer extension. Nonenzymatic template-directed RNA polymerization at the 3′-end of a primer proceeds via a bridged dinucleotide intermediate, which forms spontaneously in a pool of chemically activated nucleotides. Isocyanide nucleotide activation chemistry is incompatible with primer extension due to the required excess 2AI, which inhibits accumulation of the bridged dinucleotides.

Our laboratory has demonstrated efficient copying of various short RNA templates using 5′-phosphoro-2-aminoimidazolide-(2AI) activated ribonucleotides (2AImpN **2**) ^14^, and shown that polymerization proceeds through the *in situ* generation of 5’-5’-imidazolium-bridged dinucleotides **3** ^15^ (Fig. 1). The superiority of 2AI as a phosphate activating group over other imidazole derivatives is due at least in part to the higher accumulation and greater stability of 5’-5’-imidazolium-bridged dinucleotides ^16^. The bridged dinucleotide forms spontaneously, accumulating to a low level in a pool of chemically activated nucleotides. Activated mononucleotides hydrolyze to generate free 2AI, which in turn attacks the bridged dinucleotide to yield two 2AImpNs **2** ^16–17^. Bridged dinucleotides also decay through hydrolysis— yielding one 2AImpN **2** and one nucleoside monophosphate (NMP **1**).

A prebiotically relevant model requires *in situ* activation that is also compatible with primer extension ^11, 14, 18^. Sutherland and co-workers have recently reported a prebiotically plausible route to phosphate activation with methyl isocyanide, aldehyde, and imidazole ^19^ in a pH regime that is potentially compatible with primer extension ^16, 19^. This prompted us to seek conditions under which prebiotically plausible nucleotide activation chemistry can be applied to template-directed nonenzymatic RNA polymerization (Fig. 1).

A major hurdle to the compatibility of activation chemistry and RNA copying is that excess 2AI is required to drive nucleotide activation, but excess 2AI specifically inhibits nonenzymatic RNA polymerization by attacking the imidazolium-bridged dinucleotide intermediate (Fig. 1). We report a new pathway that both circumvents this issue and yields significantly higher concentrations of bridged dinucleotides than the spontaneous self-reaction of activated mononucleotides ^16^. Our findings point to new experiments with continuous activation of nucleotides during RNA copying.

As a first step to combining isocyanide activation with primer extension, we sought reaction conditions compatible with both. Optimal primer extension requires Mg^2+^ and mildly basic buffer (pH ~8) ^16, 20^. We examined the effects of Mg^2+^ concentration and pH on the activation of NMPs **1** to 2AImpN **2** using isocyanide (Fig. S1) and acetaldehyde. All four canonical ribonucleotides were activated under primer extension conditions (Fig. S2a). However, an undesirable full Passerini reaction product **5** ^21^, which depletes the starting NMP pool (supplementary text, Scheme S1), also formed in addition to the desired activated nucleotides (Figs. S3, Tables S1, S3). In a screen of longer chain aldehydes and ketones in place of acetaldehyde, 2-methylbutyraldehyde (2MBA) decreased the formation of **5** from 12% to 3%, while increasing the yield of 2AImpN **2** from 31% to 81% (Fig. S4, Table S2). The higher yield of **2** may stem from reduced hydrolysis of the imidoyl intermediate **4** without affecting the 2AI attack on the phosphate group.

Although the above optimizations define reaction conditions that could enable compatibility with primer extension, there remained a significant obstacle. The high concentration of 2AI required for NMP activation (Table 1) was expected to prohibit accumulation of the imidazolium bridged dinucleotides **3** necessary for RNA copying by driving the equilibrium toward 2AImpN **2** (Fig. 1) ^16^. Confirming this required a primer extension assay compatible with isocyanide activation chemistry. Because the isocyanide chemistry modified the fluorophores used for primer labelling (Fig. S5), we developed a post-labeling strategy for visualization of primer extension (supplementary text, Figs. S6, S7). Using a standard primer extension reaction in which the template sequence is 5′-CCG-3′, we found that the excess 2AI (200 mM) required for efficient activation severely inhibits primer extension in the presence or absence of activation chemistry (Fig. S8). Thus, the requirement for excess 2AI is a fundamental incompatibility between primer extension and *in situ* activation with isocyanide.

**Table 1.** Yields of 2AImpA 2 and bridged dinucleotide 3 at different 2AI concentrations measured by NMR spectroscopy.

| [2AI] mM | Activation (**2**) yield % | Bridged dinucleotide (**3**) yield % |
| --- | --- | --- |
| 10 | 6±2 | 1.8±0.5 |
| 20 | 13.4±0.6 | 2.8±0.9 |
| 50 | 28.1±0.2 | 2.1±0.2 |
| 100 | 33.0±0.5 | 0.8±0.4 |
| 200 | 48±2 | 0 |
| 400 | 34±1 | 0 |
| 800 | 19±1 | 0 |

All reactions were carried out using AMP (10 mM), acetaldehyde (200 mM), methyl isocyanide (200 mM), 2AI (varied), $Mg^{2+}$ ( $MgCl_2$ , 10 mM), and HEPES (200 mM) at pH 8.0 and $t = 6$ hours. Errors are standard deviations of the mean, $n = 3$ replicates.

Reflecting on the overall primer extension pathway— from nucleotides **1**, via activated nucleotides **2**, to the bridged dinucleotides **3** that actually promote the elongation of the primer—we asked whether the isocyanide activation chemistry might be relevant to the formation of the bridged dinucleotide as well as activated monomers. We postulated that the 2AI of pre-activated 2AImpN **2** could act as a nucleophile to displace the imidoyl leaving group of intermediate **4**, directly forming the bridged imidazolium dinucleotide **3** necessary for primer extension (Fig. 1). We therefore introduced 2AImpN **2** to a mixture of isocyanide, aldehyde, and AMP **1** without any free 2AI. We found that not only did the bridged dinucleotide species **3** form, but it accumulated to a significantly higher yield than is typical through the self-reaction of activated monomers in the absence of activation chemistry. For an equimolar mix of AMP and 2AImpA, 31P NMR spectra show 16% bridged dinucleotide **3** in the presence of isocyanide activation at t = 229 min. (the timepoint at which the concentration of bridged dinucleotide peaks), compared with only 2% in its absence (Fig. 2). To differentiate this scenario from the one in which excess 2AI drives NMP **1** activation, we call it bridge-forming activation (supplementary text).

**Figure 2.**
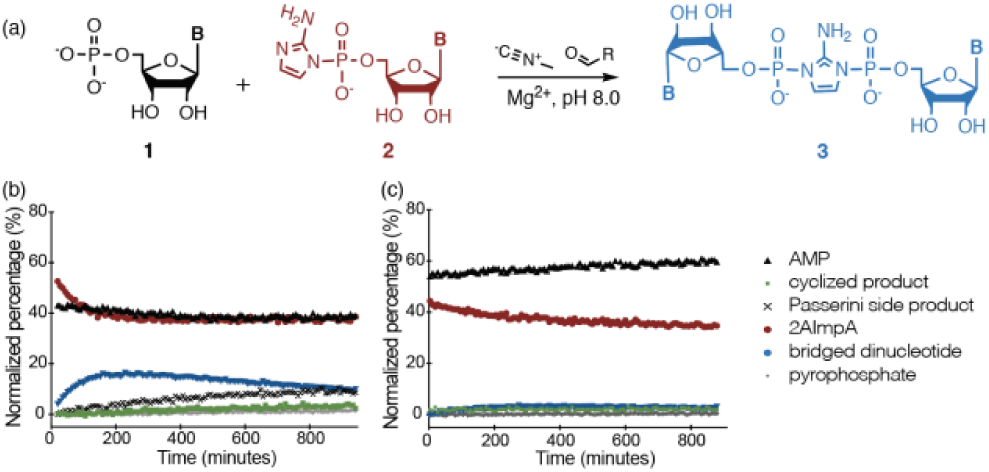
Bridge-forming activation. (a) Prebiotically plausible chemistry drives bridged dinucleotide formation. Analyses of the reaction over the course of 15 hours by 31P-NMR (b) with and (c) without bridge-forming activation. The relative percentage of each species is calculated based on the corresponding peak integration normalized to the number of phosphorus atoms. Reaction conditions: AMP (5 mM), 2AImpA (5 mM), Mg^2+^ (30 mM, MgCl_2_), HEPES (200 mM) at pH 8 with or without methyl isocyanide (200 mM) and 2MBA (200 mM).

We expect that in a prebiotically plausible scenario for RNA copying, the ratio of activated to unactivated nucleotides would vary with time. We find that bridge-forming activation functions across a broad range of ratios, with bridged dinucleotide **3** detected in every case in which activated mononucleotide **2** is present (Fig. S9a-c). Bridged dinucleotide **3** did not form in the absence of activated monomer **2** (Fig. S9d).

Interestingly, the treatment of 100% activated mononucleotides **2** with the bridge-forming activation reagents also efficiently yielded bridged dinucleotide **3** with little accompanying hydrolysis (Fig. S10): from t = 12 min. to t = 135 min., only 1% of the mononucleotides hydrolyzed whereas the bridged dinucleotide **3** yield was 40%. These observations suggest a significant contribution from a novel pathway in which the activation chemistry directly mediates the bridging of two already-activated nucleotides (rather than only the bridging of pairs of activated and unactivated nucleotides). The mechanism of this pathway is under investigation.

We next considered whether bridge-forming activation shows any preference for 2AI over 2-methylimidazole (2MI), the historically most common activating imidazole. Treatment of an equimolar mixture of AMP and 2MImpA **7** with bridge-forming activation did yield bridged dinucleotide **8**, though markedly less than with the use of 2AImpN. Without bridge-forming activation, no detectable bridged dinucleotide **8** formed (Fig. S11). The significant difference in bridged dinucleotide accumulation between 2AI-and 2MI-activated mononucleotides led us to consider how they would behave together. If bridge-forming activation is promiscuous, then treatment of an equimolar mixture of 2AI- and 2MI-activated nucleotides should yield both imidazole bridged dinucleotides. However, the reaction only yielded one detectable bridged species: 2-aminoimidazolium bridged dinucleotide **3** (Fig. S12). This demonstrates that bridge-forming activation is efficiently selective towards 2AI over 2MI in a nucleotide concentration regime that is functional in primer extension.

Encouraged by these results, we sought to apply bridge-forming activation to primer extension. To copy the 5′-GCC-3′ template, various concentrations of 2-AI activated C and G mononucleotides were mixed with unactivated C and G and treated or not treated with bridge-forming activation (Fig. 3). As a control we performed primer extension with 5 mM each 2AImpG and C and observed a distribution of +1, +2, and +3 products to serve as a baseline for other experiments (Fig. 3a). The addition of 10 mM NMPs inhibited the reaction because unactivated mononucleotides compete for the binding sites of bridged dinucleotides (compare Fig. 3a to Fig. 3b). The application of bridge-forming activation increased product yield, with 43% +3 products compared to 21% without the bridge-forming activation (Fig. 3b,c). This distribution of products from an equimolar ratio of activated and unactivated mononucleotides plus bridge-forming activation is comparable to that found with the use of 20 mM pure activated mononucleotides (Fig. 3d). Finally, applying bridge-forming activation to 10 mM pure activated mononucleotides, with no initial unactivated nucleotides, resulted in even more +3 product (53%) (Fig. 3e). Note that in these experiments the product length is template-limited, because there are no template bases beyond the +3 position. These experiments demonstrate the compatibility of isocyanide-based nucleotide activation with nonenzymatic RNA polymerization.

**Figure 3.**
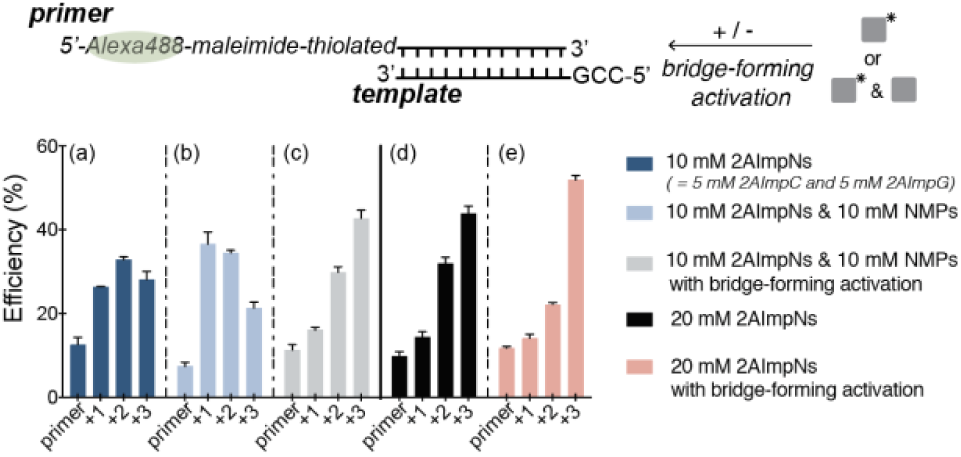
Primer extension using bridge-forming activation. Products from primer extension reactions using (a) 10 mM 2AImpN (N=C and G), (b) an equimolar mixture of 10 mM 2AImpNs and 10 mM NMPs without bridge-forming activation and (c) with bridge-forming activation, (d) 20mM 2AImpNs without bridge-forming activation and (e) with bridge-forming activation. Reaction conditions: indicated quantities of 2AImpNs and NMPs were added to primer (1 μM, green text), template (1.5 μM, black text), HEPES (200 mM) pH 8.0, and Mg^2+^ (30 mM, MgCl_2_) with isocyanide chemistry (grey and coral pink) or without (dark blue, light blue, black). Extension products were assayed by PAGE (Fig. S13) at t = 24 hours. Error bars indicate standard deviations of the mean, n = 3 replicates.

Bridge-forming activation provides several advantages for primer extension. It requires lower mononucleotide concentrations (1.5-5 mM) to generate appreciable proportions of bridged dinucleotide than required by spontaneous bridging (10-100s mM range) or direct activation with free 2AI (50 mM-400 mM NMP). The lower mononucleotide concentration combined with the higher proportion of bridged dinucleotides also raises the possibility that the Mg^2+^ concentration can be further reduced. High Mg^2+^ concentrations are notoriously problematic for primer extension, causing bridged dinucleotide hydrolysis ^22^, monomer cyclization ^20, 23–24^, and template degradation ^20, 25^. Finally, the reaction is effective over a range of unactivated to activated mononucleotide ratios (Fig. S9), a key environmental variable that would be expected to fluctuate in any plausible prebiotic scenario.

Although bridge-forming activation is compatible with primer extension and promotes the formation of the required intermediate, it depends on a source of previously activated mononucleotides. One possibility is that nucleotide activation might occur under partial dry-down conditions where all reactants including 2AI are at very high concentrations, followed by dilution to nucleotide concentrations sufficient for bridge-forming activation and primer extension, but with low enough free 2AI to minimize loss of the bridged dinucleotides. Additional processes that might sequester or degrade 2AI after efficient nucleotide activation should also be investigated. For example, UV radiation, the presumptive energy source for producing isocyanide, photodegrades 2AI on the order of days 26 although the photodegradation rates of **2** and **3** remain unknown.

Life is not expected to emerge after a single round of template-directed primer extension. A highly desirable feature of bridge-forming activation is the potential re-activation of spent nucleotides for further rounds of polymerization. Previously identified activation chemistries lead to damaging side reactions that destroy both templates and substrates ^20, 27–31^, whereas bridge-forming activation relies on RNA-compatible and specific reagents. Further work is needed to demonstrate nucleotide re-activation in the context of indefinitely continued primer extension.

## ASSOCIATED CONTENT

The Supporting Information is available free of charge on the ACS Publications website.

Supplemental Materials and Methods and Supplemental Figures and Scheme (PDF)

## AUTHOR INFORMATION

### Other Authors

**Stephanie J. Zhang** − Massachusetts General Hospital, Boston, Massachusetts, and Harvard University, Boston, Massachusetts; orcid.org/ 0000-0001-6308-5257

**Daniel Duzdevich** − Massachusetts General Hospital, Boston, Massachusetts, and Harvard Medical School, Boston, Massachusetts; orcid.org/ 0000-0002-8225-607

### Notes

The authors declare no competing financial interest.

## ACKNOWLEDGMENT

J.W.S. is an Investigator of the Howard Hughes Medical Institute. This work was supported in part by grants from the National Science Foundation (CHE-1607034) and the Simons Foundation (290363) to J.W.S. The authors thank Dr. Angelica Mariani and Dr. David Russell for their expert advice, Dr. Chun Pong Tam for assistance with methyl isocyanide synthesis, Dr. John Sutherland for sharing unpublished results and insightful conversations, Dr. Travis Walton and Ms. Lydia Pazienza for helpful discussions, and Mr. Constantin Giurgiu, Dr. Anna Wang, Dr. Travis Walton, Dr. Kyle Strom, and Dr. Zoe Todd for helpful comments on the manuscript.

## RERERENCES

(1) Szostak, J. W.; Bartel, D. P.; Luisi, P. L., Synthesizing life. Nature 2001, 409 (6818), 387–390.

(2) Wachowius, F.; Attwater, J.; Holliger, P., Nucleic acids: function and potential for abiogenesis. Q Rev Biophys 2017, 50.

(3) Lilley, D. M. J.; Eckstein, F., Ribozymes and RNA catalysis. The Royal Society of Chemistry: Cambridge, UK, 2007.

(4) Szostak, J. W., The Narrow Road to the Deep Past: In Search of the Chemistry of the Origin of Life. Angew Chem Int Edit 2017, 56 (37), 11037–11043.

(5) Horning, D. P.; Bala, S.; Chaput, J. C.; Joyce, G. F., RNA-Catalyzed Polymerization of Deoxyribose, Threose, and Arabinose Nucleic Acids. ACS Synthetic Biology 2019, 8 (5), 955–961.

(6) Joyce, G. F.; Szostak, J. W., Protocells and RNA Self-Replication. Csh Perspect Biol 2018, 10 (9).

(7) Orgel, L. E., Molecular Replication. Nature 1992, 358 (6383), 203–209.

(8) Sulston, J.; Lohrmann, R.; Orgel, L. E.; Miles, H. T., Nonenzymatic Synthesis of Oligoadenylates on a Polyuridylic Acid Template. P Natl Acad Sci USA 1968, 59 (3), 726-&.

(9) Orgel, L. E., Prebiotic chemistry and the origin of the RNA world. Crit Rev Biochem Mol 2004, 39 (2), 99–123.

(10) Weimann, B. J.; Lohrmann, R.; Orgel, L. E.; Schneide.H; Sulston, J. E., Template-Directed Synthesis with Adenosine-5’-Phosphorimidazolide. Science 1968, 161 (3839), 387-&.

(11) Prywes, N.; Blain, J. C.; Del Frate, F.; Szostak, J. W., Nonenzymatic copying of RNA templates containing all four letters is catalyzed by activated oligonucleotides. Elife 2016, 5.

(12) Vogel, S. R.; Deck, C.; Richert, C., Accelerating chemical replication steps of RNA involving activated ribonucleotides and downstream-binding elements. Chem Commun 2005, (39), 4922–4924.

(13) Westheimer, F. H., Why Nature Chose Phosphates. Science 1987, 235 (4793), 1173–1178.

(14) Li, L.; Prywes, N.; Tam, C. P.; O’Flaherty, D. K.; Lelyveld, V. S.; Izgu, E. C.; Pal, A.; Szostak, J. W., Enhanced Nonenzymatic RNA Copying with 2-Aminoimidazole Activated Nucleotides. J Am Chem Soc 2017, 139 (5), 1810–1813.

(15) Walton, T.; Szostak, J. W., A Highly Reactive Imidazolium-Bridged Dinucleotide Intermediate in Nonenzymatic RNA Primer Extension. J Am Chem Soc 2016, 138 (36), 11996–12002.

(16) Walton, T.; Szostak, J. W., A Kinetic Model of Nonenzymatic RNA Polymerization by Cytidine-5’-phosphoro-2-aminoimidazolide. Biochemistry-Us 2017, 56 (43), 5739–5747.

(17) Zhang, W.; Walton, T.; Li, L.; Szostak, J. W., Crystallographic observation of nonenzymatic RNA primer extension. Elife 2018, 7.

(18) Whitaker, D.; Powner, M. W., Prebiotic nucleic acids need space to grow. Nat Commun 2018, 9.

(19) Mariani, A.; Russell, D. A.; Javelle, T.; Sutherland, J. D., A Light-Releasable Potentially Prebiotic Nucleotide Activating Agent. J Am Chem Soc 2018, 140 (28), 8657–8661.

(20) Szostak, J. W., The eightfold path to non-enzymatic RNA replication. Journal of Systems Chemistry 2012, 3 (1), 2.

(21) Sutherland, J. D.; Mullen, L. B.; Buchet, F. F., Potentially prebiotic Passerini-type reactions of phosphates. Synlett 2008, (14), 2161–2163.

(22) Walton, T.; Pazienza, L.; Szostak, J. W., Template-Directed Catalysis of a Multistep Reaction Pathway for Nonenzymatic RNA Primer Extension. Biochemistry-Us 2019, 58 (6), 755–762.

(23) Breslow, R.; Huang, D. L., Effects of Metal-Ions, Including Mg2+ and Lanthanides, on the Cleavage of Ribonucleotides and Rna Model Compounds. P Natl Acad Sci USA 1991, 88 (10), 4080–4083.

(24) Harada, K.; Orgel, L. E., The Cyclization of Arabinosyladenine-5’-Phosphorimidazolide. J Mol Evol 1991, 32 (5), 358–359.

(25) Adamala, K.; Szostak, J. W., Nonenzymatic Template-Directed RNA Synthesis Inside Model Protocells. Science 2013, 342 (6162), 1098–1100.

(26) Todd, Z. R.; Szabla, R.; Szostak, J. W.; Sasselov, D. D., UV photostability of three 2-aminoazoles with key roles in prebiotic chemistry on the early earth. Chem Commun (Camb) 2019, 55 (70), 10388–10391.

(27) Biron, J. P.; Pascal, R., Amino acid N-carboxyanhydrides: Activated peptide monomers behaving as phosphate-activating agents in aqueous solution. J Am Chem Soc 2004, 126 (30), 9198–9199.

(28) Powner, M. W.; Gerland, B.; Sutherland, J. D., Synthesis of activated pyrimidine ribonucleotides in prebiotically plausible conditions. Nature 2009, 459 (7244), 239–242.

(29) Leman, L.; Orgel, L.; Ghadiri, M. R., Carbonyl sulfide-mediated prebiotic formation of peptides. Science 2004, 306 (5694), 283–286.

(30) Leman, L.; Orgel, L.; Ghadiri, M. R., Amino acid condensation mediated by carbonyl sulfide. Abstr Pap Am Chem S 2004, 228, U693–U693.

(31) Ferris, J. P.; Hagan, W. J., Hcn and Chemical Evolution - the Possible Role of Cyano Compounds in Prebiotic Synthesis. Tetrahedron 1984, 40 (7), 1093–1120.

