## Supplementary Material for "Prebiotically Plausible Activation Chemistry Compatible with Nonenzymatic RNA Copying"

### Contents

**1. Materials and Methods** . . . . . **S3**
1.1. General information . . . . . S3
1.2. Synthesis and storage of methyl isocyanide . . . . . S3
1.3. Preparation, storage, and concentration determination of stock solutions . . . . . S3
1.4. Nucleoside 5′-phosphoro-2-aminoimidazolides (2AImpN) . . . . . S3
1.4.1. Generation by isocyanide nucleotide activation chemistry . . . . . S3
1.4.2. Synthesis of synthetic standard . . . . . S4
1.5. Synthesis of adenosine 5′-phosphoro-2-methylimidazolides (2MImpA) . . . . . S4
1.6. Primer extension reaction and analysis of the polymerization products . . . . . S4
**2. Supplementary Figures** . . . . . **S5**
**3. Supplementary Scheme** . . . . . **S19**
**4. Supplementary Table** . . . . . **S20**
**5. Supplementary Text** . . . . . **S22**
**6. Supplementary References** . . . . . **S23**

### 1. General information

*Materials.* Reagents and solvents were obtained with highest purity available from Acros Organics, Alfa Aesar, Fisher Scientific, Sigma-Aldrich, ThermoFisher Scientific, or Tokyo Chemical Industry Co., and were used without any further purification unless noted below. Nucleoside-5′-monophosphate, free acid, was purchased from Santa Cruz Biotechnology. 2-aminoimidazole hydrochloride and 2,2′-dipyridyldisulfide were purchased from Combi Blocks. RNA oligonucleotides were purchased from Integrated DNA Technologies. All reactions were carried out in DNase/RNase-free distilled water.
*NMR spectroscopy.* $^{1}H$ and $^{31}P$-NMR spectra were obtained using a 400 MHz NMR spectrometer (Varian INOVA) operating at 400 MHz and 161 MHz respectively. Samples in $H_2O/D_2O$ mixtures were analyzed using Wet1D suppression to collect $^{1}H$-NMR data. Chemical shifts (δ) are shown in ppm. Coupling constants (J) are given in Hertz (Hz) and the notations s, d, t, and m represent the multiplicities of singlet, doublet, triplet, and multiplet, respectively.
*pH measurements.* pH values were determined by a micro pH probe (Orion 9863BN) equipped with a needle tip and a SevenCompact meter (Mettler Toledo S220).
*Mass spectrometry.* All samples of nucleotides from isocyanide activation reactions were purified by acetone precipitation as described previously, prior to analysis by mass spectrometry. [1] The nucleotides were precipitated by combining the activation reaction mixture with a solution of 3 mL diethyl ether, 6 mL of acetone, and 0.23 g of sodium perchlorate. The precipitated material was washed with acetone and diluted to 200 μM in Milli-Q water with a few drops of acetonitrile immediately prior to analysis. Spectra were obtained by direct injection on an Esquire 6000 mass spectrometer (Bruker Daltonics), operated in the alternating ion mode.
*Data analysis.* All spectra were analyzed using MestReNova (version 12.0.3). The yields of conversion were determined by the relative integration of the signals in the $^{1}H$ or $^{31}P$ NMR spectra.
All data shown are representative of distinct samples, n = 3 replicates or greater.

### 2. Synthesis and storage of methyl isocyanide [2]

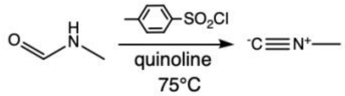


In a 500 mL 3-neck flask equipped with a pressure-equalizing dropping funnel, a sealed mechanical stirrer, a thermometer, and a receiver trap were placed 100 mL (800 mmoles) of quinoline, which was freshly distilled from zinc dust (Alfa Aesar), and 57.2 g (300 mmoles) of *p*-toluenesulfonyl chloride. The solution was heated to 75 °C in an oil bath and the system evacuated to a pressure of 15 mm. The receiver was cooled in a bath of liquid nitrogen. While the solution was vigorously stirred and maintained at this temperature, 11.8 g (200 mmoles) of N-methylformamide was added dropwise to maintain a smooth distillation rate. The addition was complete in 45-60 minutes. The material, which collected in the receiver, was distilled under vacuum. Methyl isocyanide was collected at 59-60°C.

Analysis by NMR indicates that the purity exceeded 98% (Fig. S1); $^{1}H$-NMR (400MHz, 10% $D_2O$ in $H_2O$) δ 3.03 (t, $^{1}H$, J =2.28 Hz). Methyl isocyanide was stored at pH 9 or greater directly after distillation in Teflon-backed screw-cap glass vials inside air-tight plastic containers.

### 3. Preparation, storage, and concentration determination of stock solutions

Stock solutions of nucleoside-5′-monophosphate disodium salt, 2-aminoimidazole hydrochloride, and buffer, 2-(Bis (2-hydroxyethyl) amino) acetic acid (BICINE), were prepared by dissolving the corresponding reagent in DNase and RNase-free distilled water. After adjusting the pH to the reported values with NaOH/HCl, the stock solution was filter sterilized with 0.22-micron syringe filters (Millipore Sigma). Each stock solution was then aliquoted and kept at -20 °C until further use.

The exact concentrations of the nucleoside-5′-monophosphates were determined by analysis of serial dilutions on a spectrophotometer. The absolute concentrations of the other stock solutions were found by comparing the integrals of $^{1}H$-NMR peaks of interest to the calibrant, adenosine-5′-monophosphate by NMR spectroscopy.

### 4. Nucleoside 5′-phosphoro-2-aminoimidazolides (2AImpN)

#### 1.4.1. Generation by isocyanide nucleotide activation chemistry

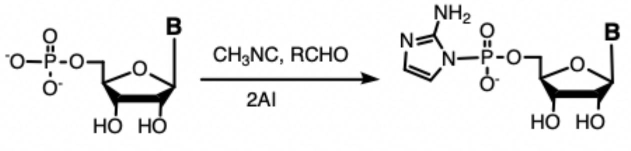


1 2

The nucleoside-5′-monophosphate (NMP), 2-aminoimidazole (2AI), the indicated aldehyde, magnesium chloride ($MgCl_2$), water, and/or buffer solution were added to their corresponding concentration listed in Table S1, in a total volume of 450 μL. Buffer was added to avoid changing pH of the solution during the reaction. Methyl isocyanide was added to the solution, which was briefly vortexed ensuring proper mixing. The total volume was brought to 450 μL. The reaction was allowed to sit for 6 hours, which was the optimal incubation time as determined by the time course (Fig. 2), at room temperature. 50 μL $D_2O$ (10%) was added before transferring 500 μL of this mixture to an NMR tube (Fisher) for NMR spectroscopy. The chemical shifts of 2AImpA were reported as they appeared in the $^{31}P$ NMR of the reaction mixture. Spectroscopic data (Fig. 3a & b) were in agreement with previous literature [3-4]; for 2AImpA, $^{31}P$-NMR (161 MHz, 10% $D_2O$ in $H_2O$, 1H-decoupled): δ -8.70.

The progress of the reaction was monitored by $^{31}P$-NMR over the course of 15 hours in a Shigemi NMR microtube assembly (Sigma Aldrich) with trimethyl phosphate as the internal reference (δ 0.00 ppm). The entire time-course of the reaction was obtained through arrayed acquisition and tracked by monitoring peak areas.

#### 1.4.2. Synthesis of synthetic standards for activated nucleotides

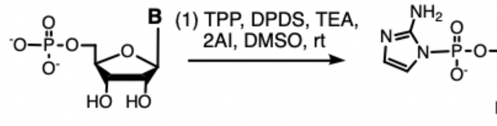


Nucleoside-5′-monophosphate free acid (1 equiv.) was dissolved along with 2-aminoimidazole hydrochloride (10 equiv.) in water and the pH was adjusted to 5.5 using NaOH/HCl. The solution was flash frozen in liquid nitrogen and lyophilized to yield a powder. To the solution of nucleoside-5′-mono-phosphate and 2-aminoimidazole in triethylamine (13 equiv., TEA) and DMSO (30 mL) was added 2,2′-dipyridyldisulfide (10 equiv., DPDS), triphenylphosphine (9 equiv., TPP) with stirring and under argon for 30 minutes. The reaction mixture was then added to a pre-chilled solution of acetone/diethyl ether/TEA (400/250/30 mL) to which 1.5mL of saturated $NaClO_4$ in acetone was added. After the precipitate had settled out, the majority of the supernatant was removed using pipette-suction and the remaining suspension was centrifuged at 4,000 rpm for 5 min. The pellets were washed first with a solution of acetone/diethyl ether/TEA (133/83/10 mL) and twice with acetone (10 mL). The product was purified by reverse-phase flash chromatography using gradient elution between (A) aqueous Milli-Q water and (B) acetonitrile. The sample was eluted between 0% and 15% B over 8 column volumes (CVs) with a flow rate of 40 mL/min.

### 5. Synthesis of adenosine 5′-phosphoro-2-methylimidazolides (2MImpA)

2MImpA was prepared as above but with the following minor difference: 2-methylimidazole was used instead of 2-aminoimidazole.

### 6. Primer extension reactions and analysis of polymerization products

The primer-template duplex was first annealed in a 40 μL solution containing 2 μM thiol-modified primer, 3.1 μM template, 416.7 mM HEPES (pH 8.0), 62.5 mM $MgCl_2$ by heating at 95 °C for 3 min. and cooling down to 23 °C at a rate of 0.1 °C/s. The reaction was initiated by the addition of activated monomers. The stock solutions (100 mM) of 2AI-activated monomers had a pH of around 9.6.

Aliquots (10 μL each) were removed at given time points and desalted using a ZYMO Oligo Clean & Concentrator spin column (ZYMO Research). The isolated material was resuspended in 30 uL 100 mM HEPES (pH 7.50) and disulfide bonds were reduced using a 10-fold molar excess of tris-(2-carboxyethyl) phosphine hydrochloride (TCEP). Alexa 488 C5 maleimide dissolved in anhydrous dimethyl sulfoxide (DMSO) at a concentration of 1 mM was added to the primer-template duplex dropwise, and the reaction was allowed to proceed at room temperature for 2 hours, protected from light. The labeled primer-template duplex was separated from free dye using ZYMO DNA Clean & Concentrator-5 spin columns, resuspended in 5 uL of 100 mM HEPES buffer, and mixed with 30 μL of quenching buffer containing 8.3 M urea, 1.3x Tris/Borate/EDTA (TBE) buffer (pH 8.0), 0.005% Bromphenol Blue, 0.04% Orange G, and 75 μM RNA complementary to the template. To denature the labeled product from the template prior to polyacrylamide gel analysis, and sequester the template with its complement (added in excess as a competitor), the sample was heated to 95 °C for 3 min. and cooled to 23 °C at 0.1 °C/s. The primers, templates, complementary DNAs and RNAs used in the primer extension assays are listed below (Table S4).

20% polyacrylamide gels were prepared using the SequaGel–UreaGel system (National Diagnostics, Atlanta, GA). The gels were allowed to pre-run at a constant power of 20 W for 40−45 mins. 15 μL aliquots of samples were separated by 20% (19:1) denaturing PAGE at a constant power of 5 W for 10 min. and 25 W for 1.5 hrs. Gels were imaged on a Typhoon 9410 scanner (GE Healthcare, Little Chalfont, Buckinghamshire, UK) and quantified using the accompanying ImageQuant TL software. All reactions were performed in triplicate or greater.

### 2. Supplementary Figures

**Figure S1.** $^1$H-NMR spectrum of synthesized methyl isocyanide in 10% $D_2O$. Inset is the zoomed-in view of the three peaks.

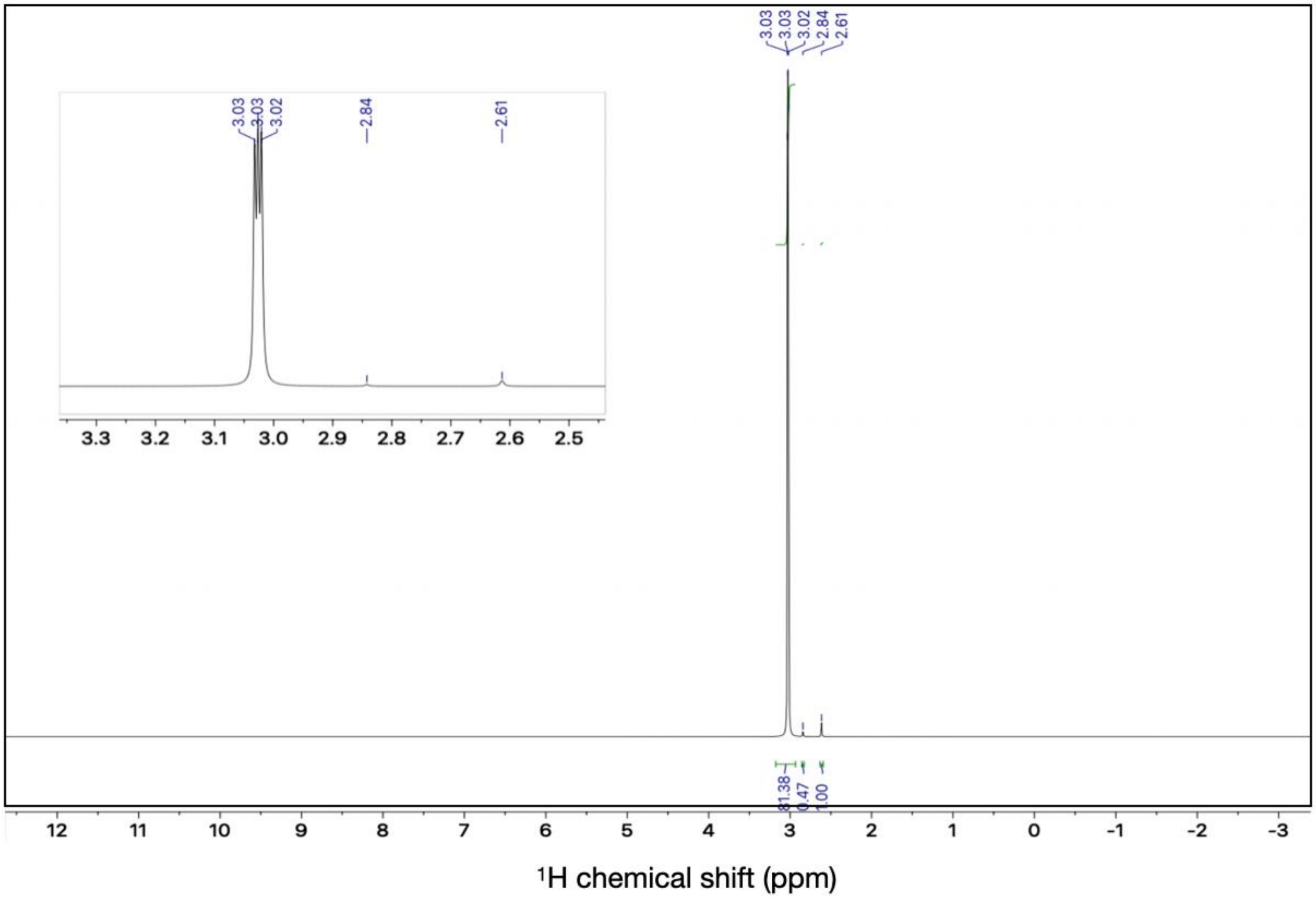

**Figure S2a. Isocyanide-mediated phosphate activation with 2AI of the four canonical ribonucleotides.** The relative yield of each 2AImpN is calculated based on the corresponding $^{31}$P NMR peak integration normalized to the number of phosphorus atoms in a given species. Errors are standard deviations of the mean, n = 3 replicates.
Reaction conditions: AMP (10 mM), $Mg^{2+}$ (30 mM, $MgCl_2$), methyl isocyanide (200 mM), acetaldehyde (200 mM), 2AI (100 mM), and HEPES (200 mM) at pH 8 and t = 6 hours (the timepoint at which the concentration of activated monomer plateaus) (Fig. S2b).

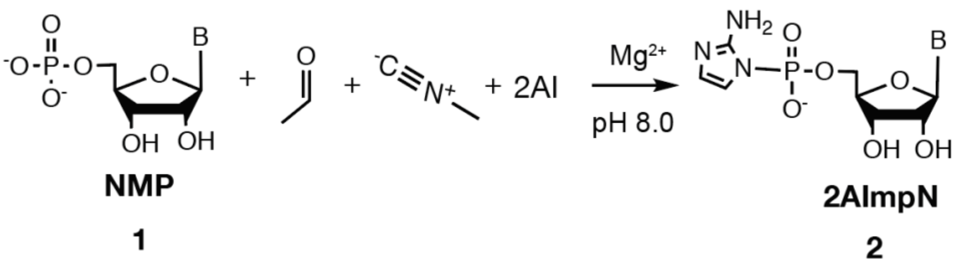


**2AImpG** (39±3%)

**2AImpC** (40±3%)

**2AImpA** (36±3%)

**2AImpU** (30±2%)

**Figure S2b. $^{31}$P-NMR time-course experiments for 2AImpA synthesis by isocyanide chemistry.** (a) The stacked $^{31}$P-NMR spectra of nucleotide activation. All spectra are referenced to the trimethyl phosphate peak at 0 ppm (not shown). (b) Quantification of nucleotide activation as a function of time based on (a).
Reaction conditions: AMP (10 mM), $Mg^{2+}$ (10 mM, $MgCl_2$), HEPES (200 mM) at pH 8, 2AI (200 mM), acetaldehyde (200 mM), and methyl isocyanide (200 mM).

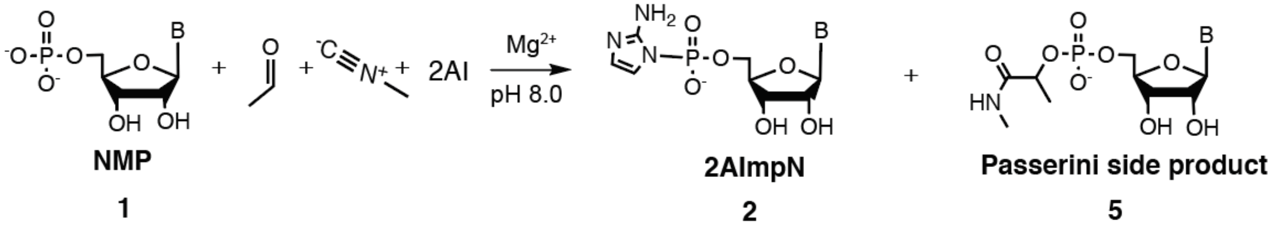


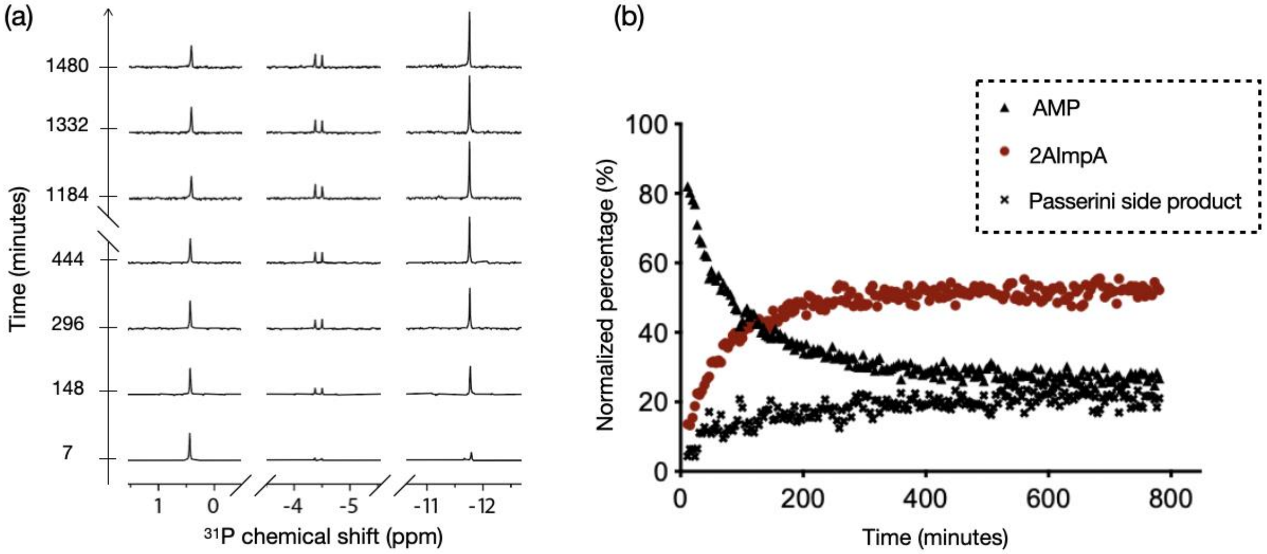

**Figure S3. Formation of 2AImpA by isocyanide activation chemistry.** $^{31}$P NMR spectra of (a) AMP alone. (b) 2AImpA formed by isocyanide activation chemistry (c) as in (b) but with spiking of synthetic standard 2AImpA.
Initial reaction conditions:
(a) AMP (10 mM), $Mg^{2+}$ (30 mM, $MgCl_2$), and HEPES (200 mM) at pH=8.0.
(b) AMP (10 mM), acetaldehyde (150 mM), methyl isocyanide (200 mM), $Mg^{2+}$ (30 mM, $MgCl_2$), HEPES (200 mM) at pH 8, 2AI (100 mM).

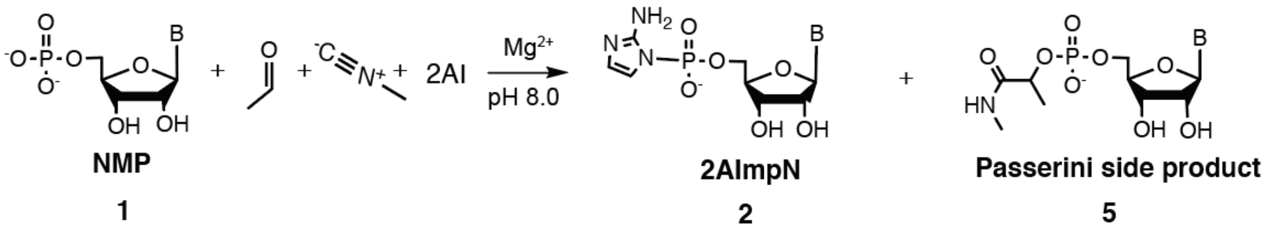


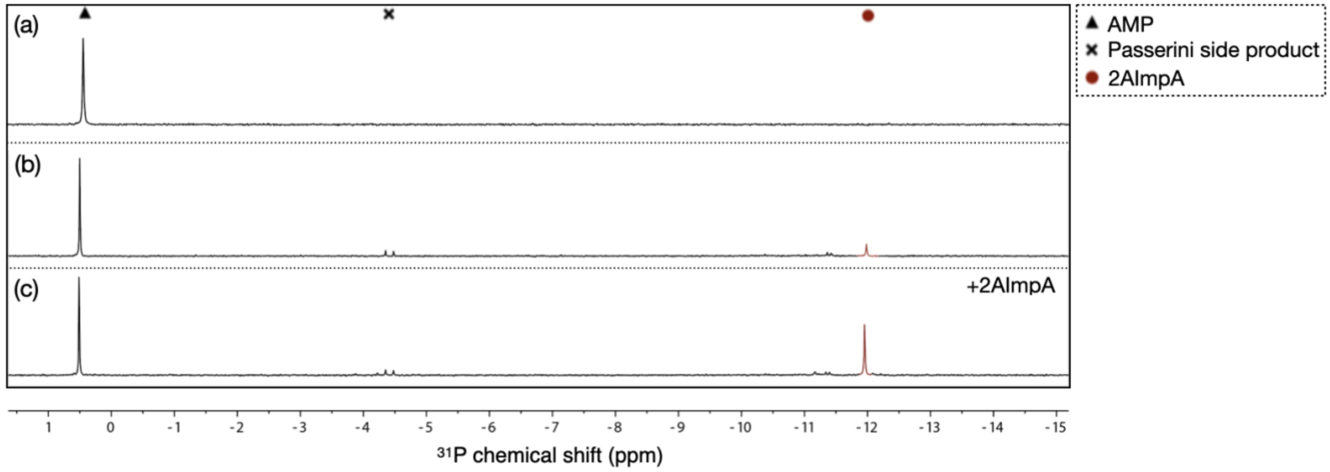

**Figure S4. 2-methyl butyraldehyde (2MBA) yields efficient activation with minimal Passerini side product.** A $^{31}$P-NMR spectrum of the standard activation reaction mixture in 10% $D_2O$ in $H_2O$ at t = 6 hr.
Reaction conditions: AMP (10 mM), 2MBA (200 mM), methyl isocyanide (200 mM), $Mg^{2+}$ (30 mM, $MgCl_2$), HEPES (200 mM) at pH 8, and 2AI (100 mM).

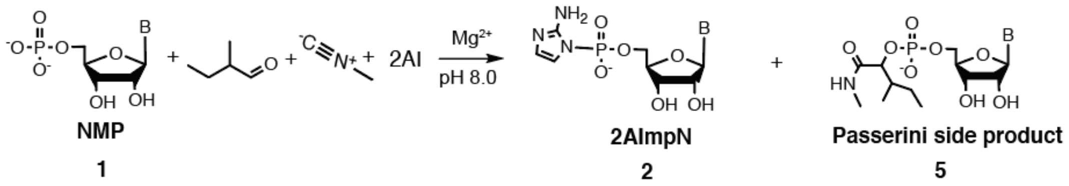


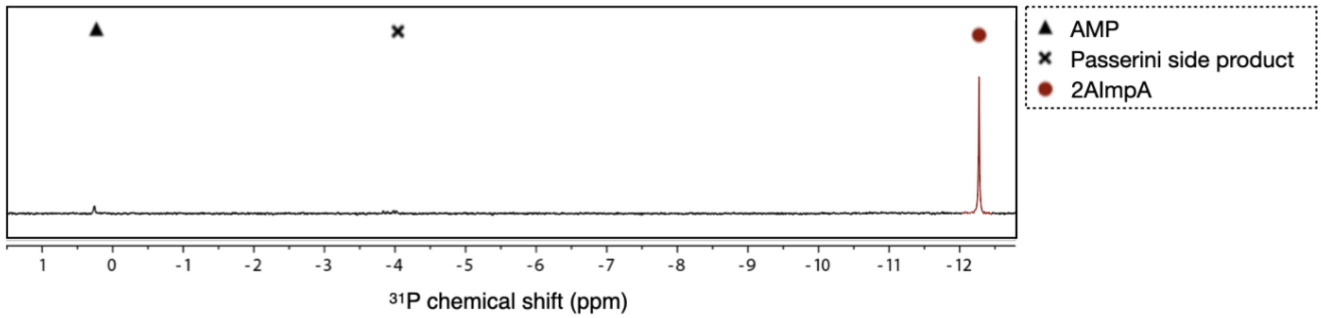

**Figure S5.** Instability of dye-labeled primers to isocyanide nucleotide activation chemistry. Fluorescent dyes expected to be the least likely to react with the isocyanide nucleotide activation chemistry and within the relevant imaging wavelength range were tested. Most dyes contain either carboxyl and/or amine functional groups. We tested dyes that contain only hindered tertiary amines, which are less likely to participate in the chemistry. However, PAGE analysis revealed additional bands, band shifts, and band disappearances.

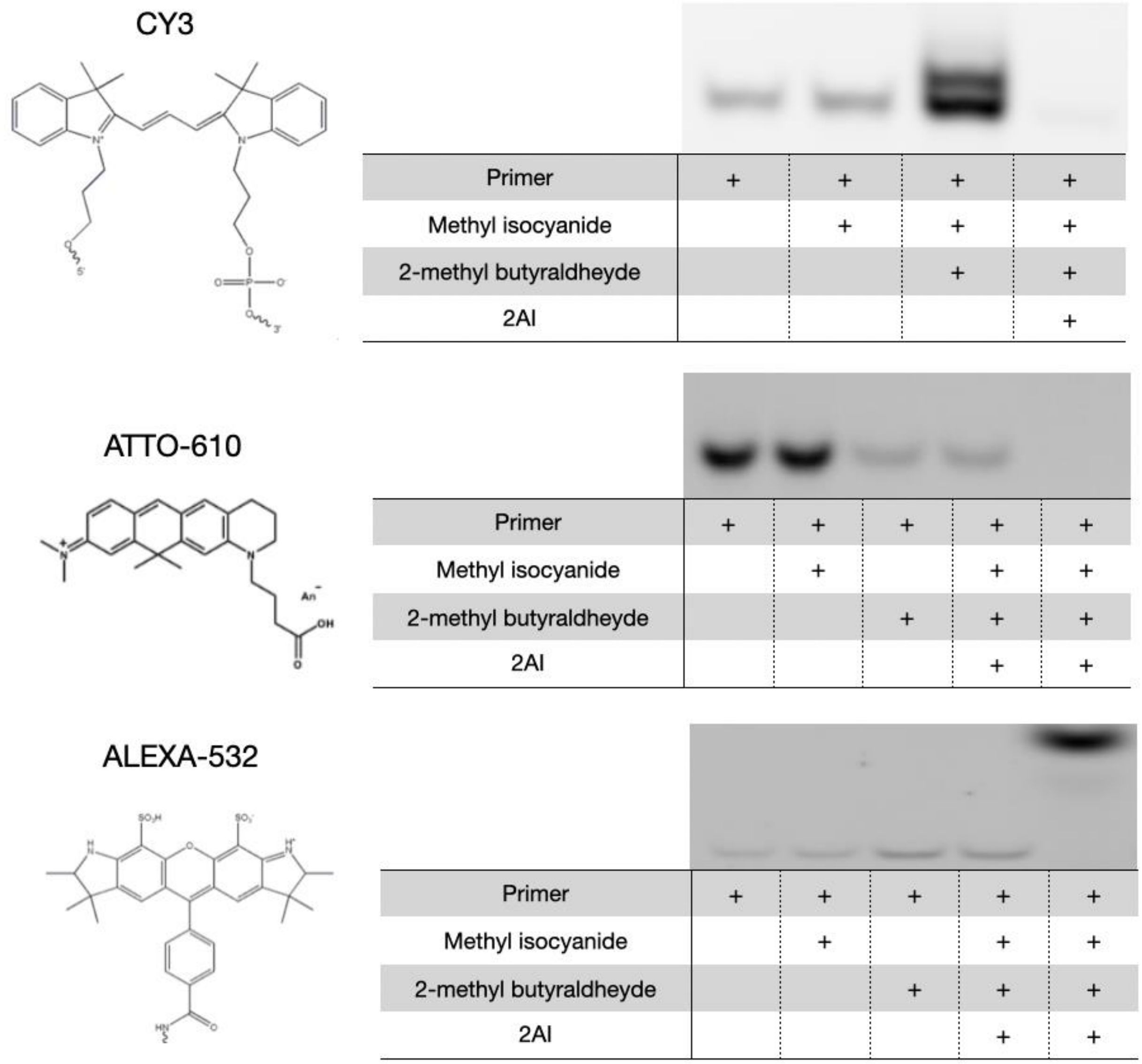


| DYE | OBSERVATIONS |
|---|---|
| CY3 | An additional band was observed when the primers were treated with isocyanide and aldehyde. |
| ATTO-610 | The disappearance of the primer band was observed on treatment with isocyanide and aldehyde, with or without 2AI. |
| ALEXA-532 | A band shift was observed when the primers were treated with isocyanide, aldehyde, and 2AI. |

**Figure S6.** Schematic representation of post-labeling used to visualize the primer extension. From left to right, each primer was treated with: (1) nothing, (2) methyl isocyanide, (3) 2MBA, (4) 2AI, (5) methyl isocyanide and 2MBA, (6) methyl isocyanide, 2MBA, and 2AI. Three trials are shown. The band intensity varies from trial to trial, probably due to variability in purification and labeling efficiency, which should not affect internally normalized quantifications of primer extension efficiencies.

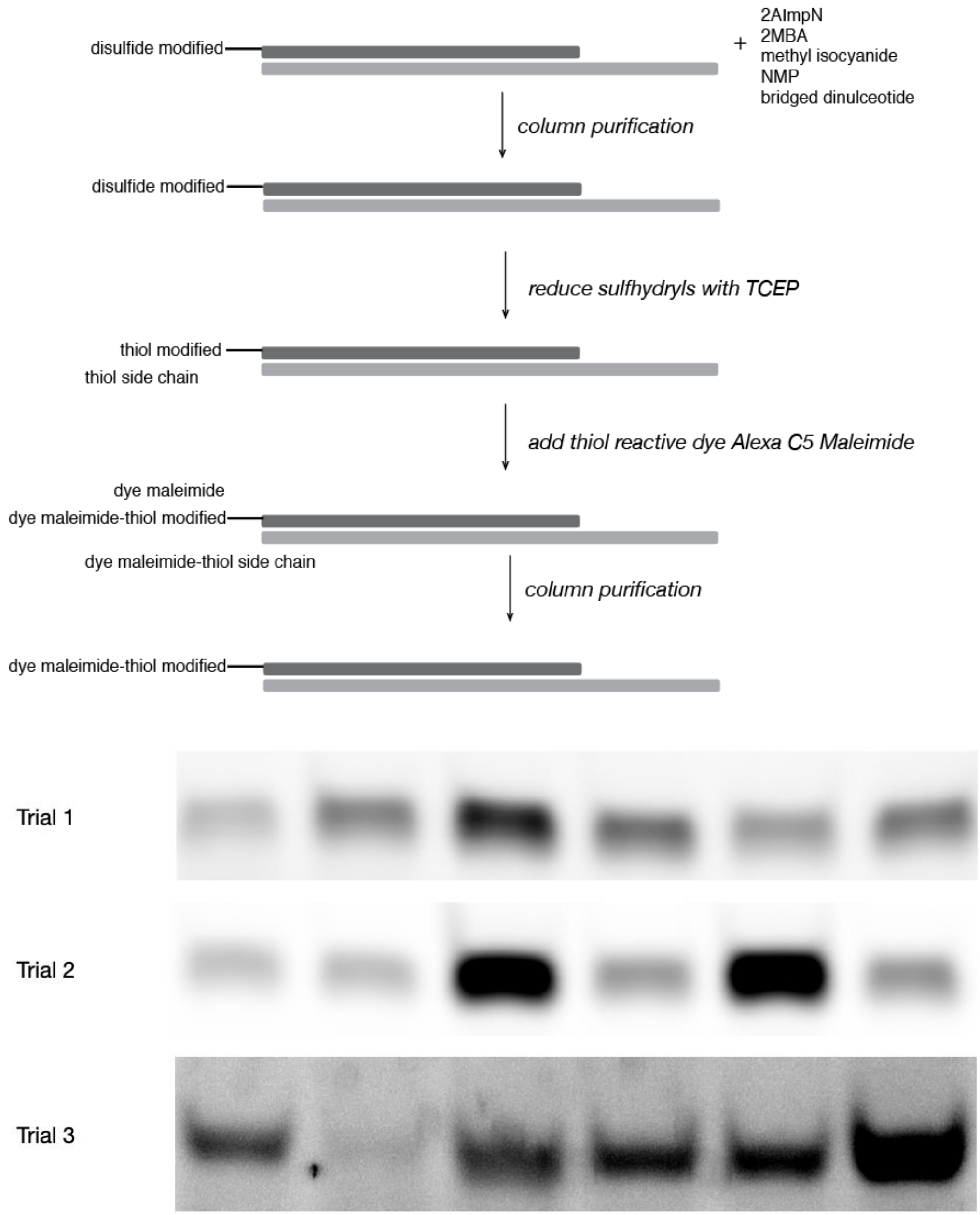


| Primer | + | + | + | + | + | + |
|---|---|---|---|---|---|---|
| Methyl isocyanide | | + | | | + | + |
| 2-Methyl butyraldehyde | | | + | | + | + |
| 2AI | | | | + | | + |

**Figure S7.** Primer extension efficiency measured by pre-labeling or post-labeling methods in the absence of isocyanide chemistry. Purified activated monomers were added to (a) the pre-labeled and (b) thiol-modified primer (triplicate data shown in both cases). A quantification of the results across the two methods (c) shows very good agreement. The variance in intensity seen in (b) is consistent with Figure S8.

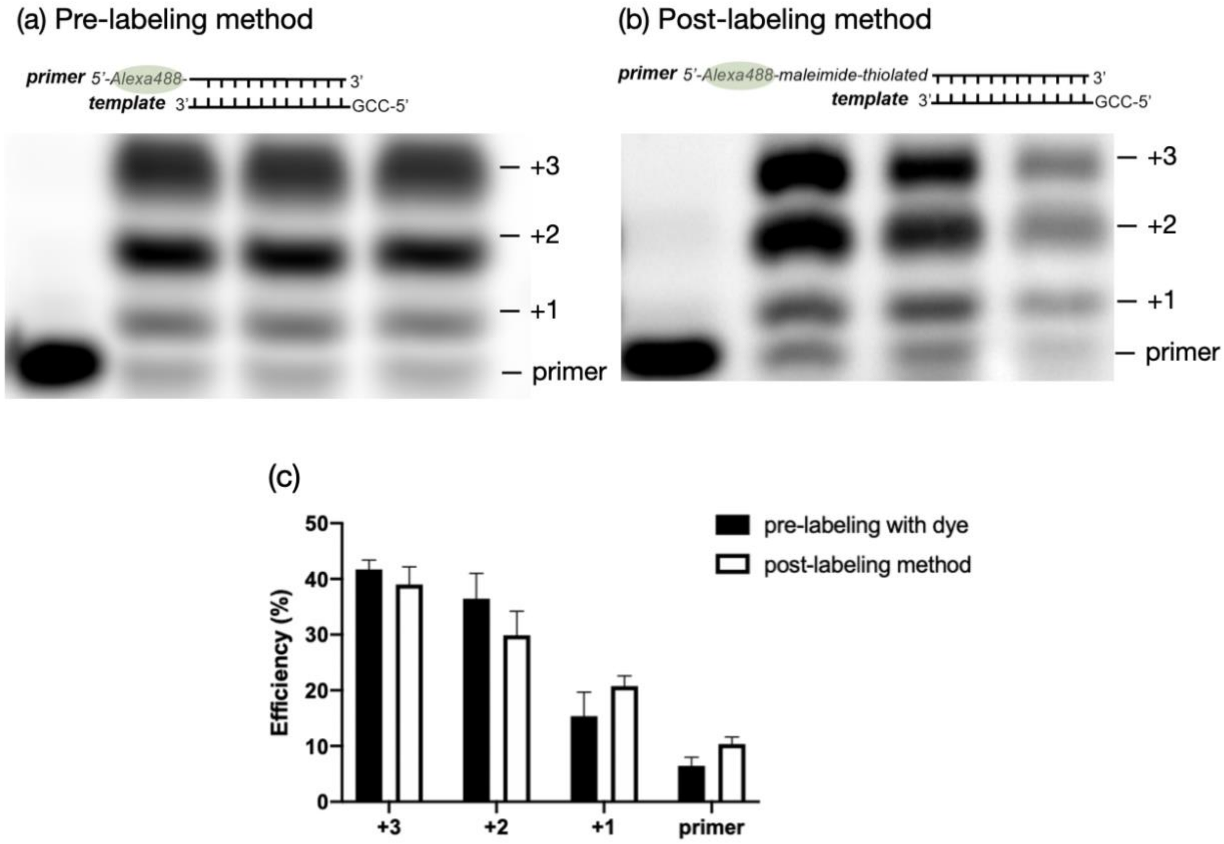

**Figure S8.** Excess 2AI inhibits primer extension. In the presence of 200 mM 2AI, only minimal +1 product was observed in reactions with activated mononucleotides, or unactivated mononucleotides with isocyanide chemistry (lanes 5-8). This is in contrast to the positive control (lanes 2-4), where primers were extended to +3 in the absence of added 2AI.

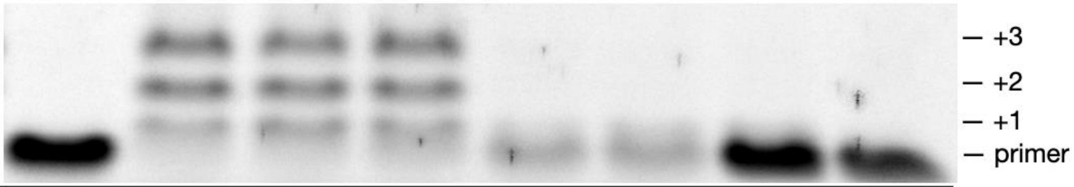


| | 1 | 2 | 3 | 4 | 5 | 6 | 7 | 8 |
|---|---|---|---|---|---|---|---|---|
| 2AImpC&G (mM) | - | 10 each | 10 each | 10 each | 10 each | 10 each | - | - |
| CMP&GMP (mM) | - | - | - | - | - | - | 10 each | 10 each |
| 2AI (mM) | - | - | - | - | 200 | 200 | 200 | 200 |
| Isocyanide chemistry | - | - | - | - | - | - | + | + |

**Figure S9. Isocyanide chemistry drives bridged dinucleotide formation with various ratios of NMP to 2AImpN.** Bridged dinucleotides can be generated across various ratios of 2AImpA to AMP: (a) ~15:85, (b) ~50:50, (c) ~80:20. Analyses of the reactions over the course of 15 hours; insets are from the same data, highlighting the bridged dinucleotides. (d) No bridged dinucleotide in the absence of activated monomer at the ratio of 2AImpA to AMP at 0:100. The relative percentage of each species is calculated based on the corresponding peak integration normalized to the number of phosphorus atoms. Passerini side product formation exhibits no correlation with the ratio of 2AImpA to AMP, although 5% more of it forms than in the absence of activated monomer because the reaction proceeds via the imidoyl phosphate **4** (Scheme S1), which undergoes competing intramolecular attack by hydroxyl to form **5**.
Reaction conditions: varying ratio of 2AImpA/AMP (10 mM total nucleotides), $Mg^{2+}$ (30 mM, $MgCl_2$), HEPES (200 mM) at pH 8 with or without methyl isocyanide (200 mM), and 2MBA (200 mM).

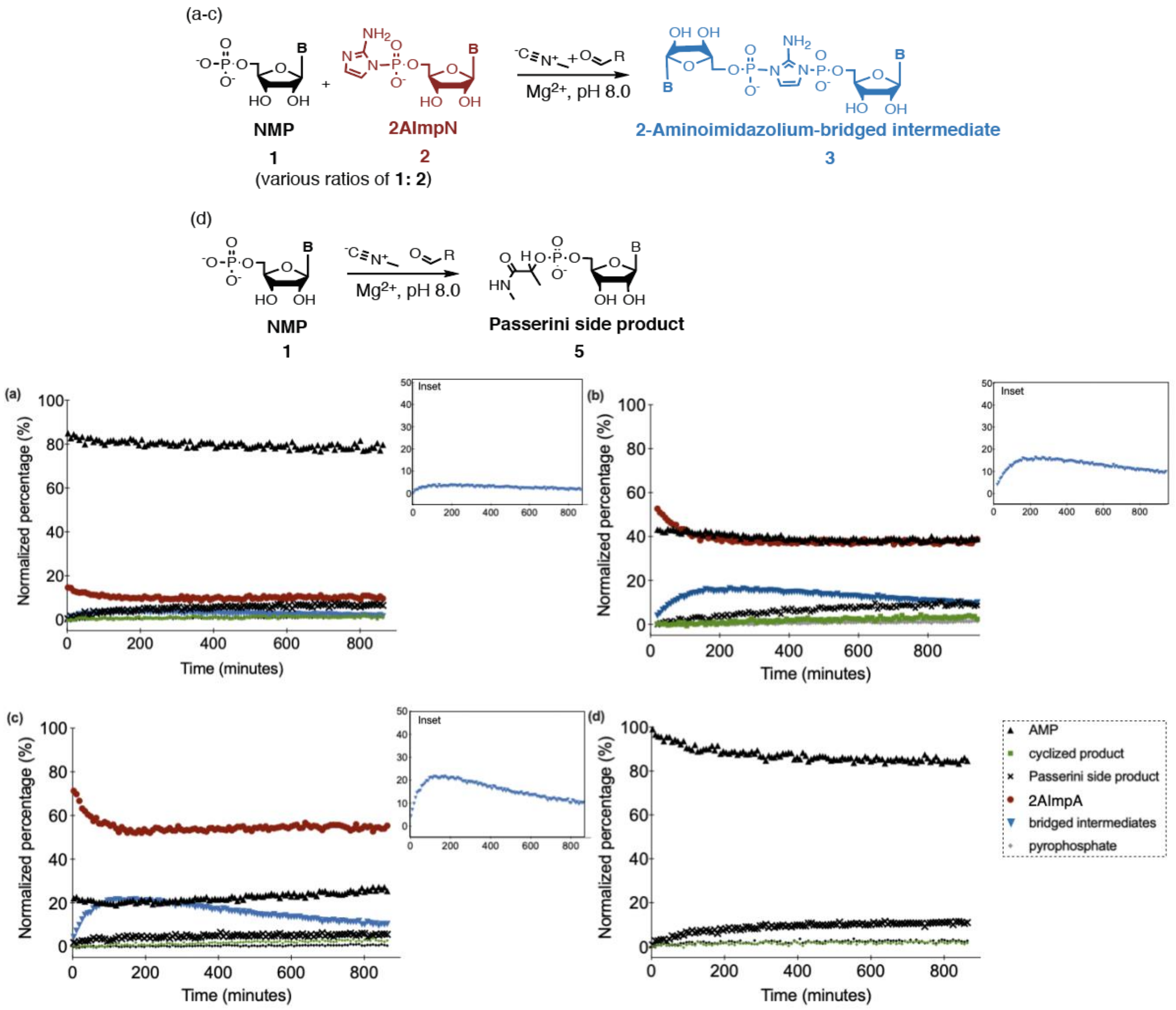

**Figure S10. Isocyanide chemistry drives bridged dinucleotide formation even with only 2AI-activated mononucleotides as substrates.** (a) The prebiotically plausible synthesis of bridged dinucleotide using only 2AI-activated monomers. Analyses of the reaction over the course of 15 hours (b) without and (c) with bridge-forming chemistry. The relative percentage of each species is calculated based on the corresponding peak integration normalized to the number of phosphorus atoms.
Reaction conditions: 2AImpA (10 mM), $Mg^{2+}$ (30 mM, $MgCl_2$), HEPES (200 mM) at pH 8 with or without methyl isocyanide (200 mM), and 2MBA (200 mM).

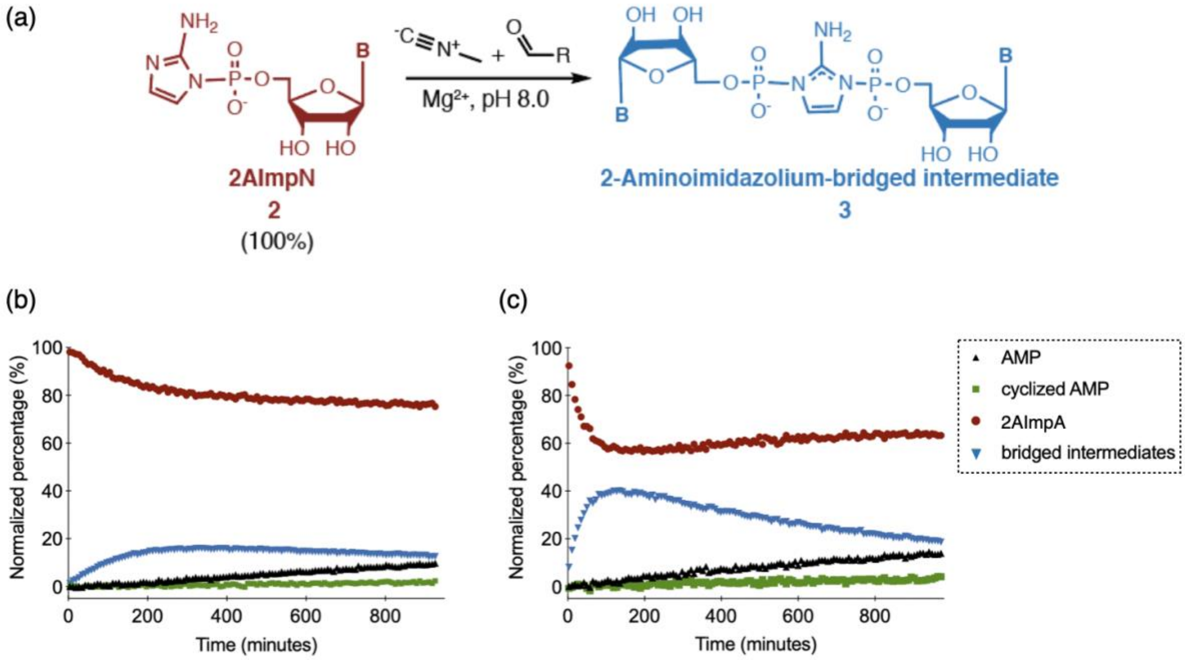

**Figure S11.** Bridge-forming chemistry acting on an equimolar mixture of AMP and 2MImpA. (a) Scheme for synthesis of bridged intermediates from a mixture of AMP and 2MImpA. Analyses of the reaction over the course of 15 hours (b) without and (c) with bridge-forming chemistry. The relative percentage of each species is calculated based on the corresponding peak integration normalized to the number of phosphorus atoms. (Much higher concentrations of activated mononucleotides are required to spontaneously form the bridged dinucleotide with 2MI, which explains the lack of bridged dinucleotide in the control case. [5])
Reaction conditions: 2MImpA (5 mM), AMP (5 mM), $Mg^{2+}$ (30 mM, $MgCl_2$), HEPES (200 mM) at pH 8 with or without methyl isocyanide (200 mM), and 2MBA (200 mM).

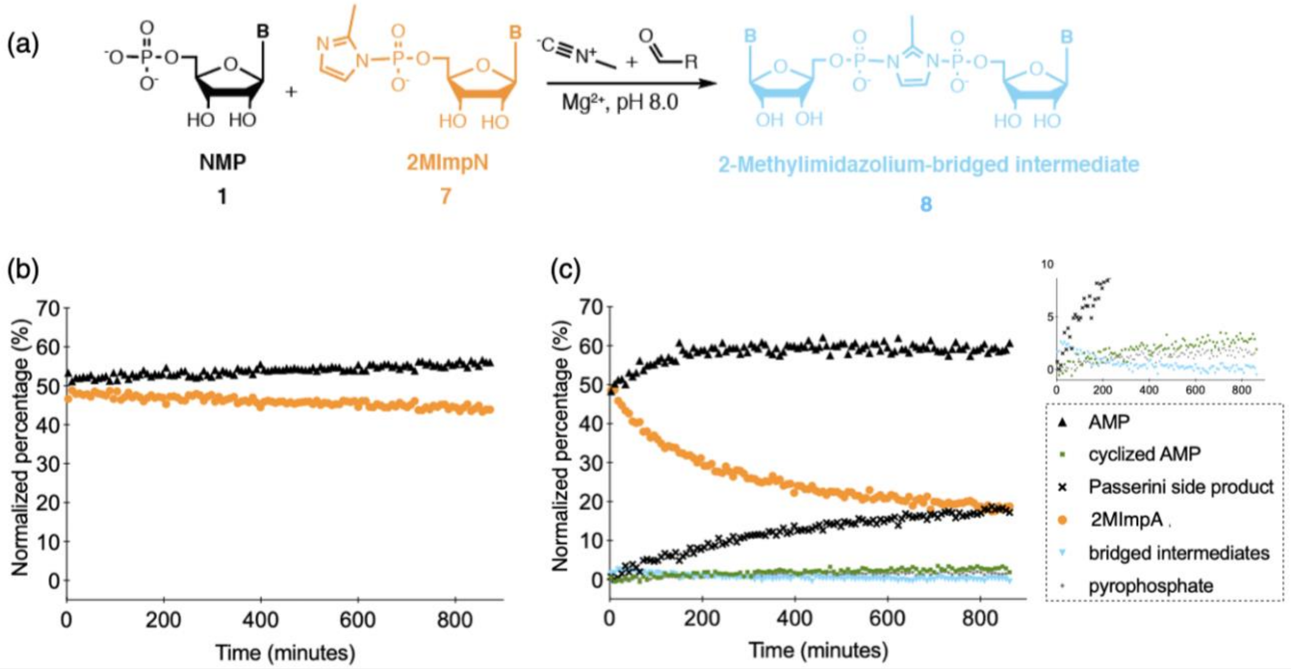

**Figure S12.** The selectivity of bridge-forming chemistry towards 2-aminoimidazolium bridged dinucleotides. (a) Scheme for synthesis of bridged dinucleotide using an equimolar mixture of 2AImpA and 2MImpA. Analyses of the reaction over the course of 15 hours (b) without and (c) with bridge-forming chemistry. The relative percentage of each species is calculated based on the corresponding peak integration normalized to the number of phosphorus atoms. (d) The only bridged species (δ -12.95) present in the synthesis can be assigned to 2AImpA because the shift of 2-aminoimidazolium bridged dinucleotide is at δ -13.0, whereas 2-methylimidazolium bridged dinucleotide is at δ -14.02.
Reaction conditions: 2AImpA (5 mM), 2MImpA (5 mM), $Mg^{2+}$ (30 mM, $MgCl_2$), HEPES (200 mM) at pH 8 with or without methyl isocyanide (200 mM), 2MBA (200 mM).

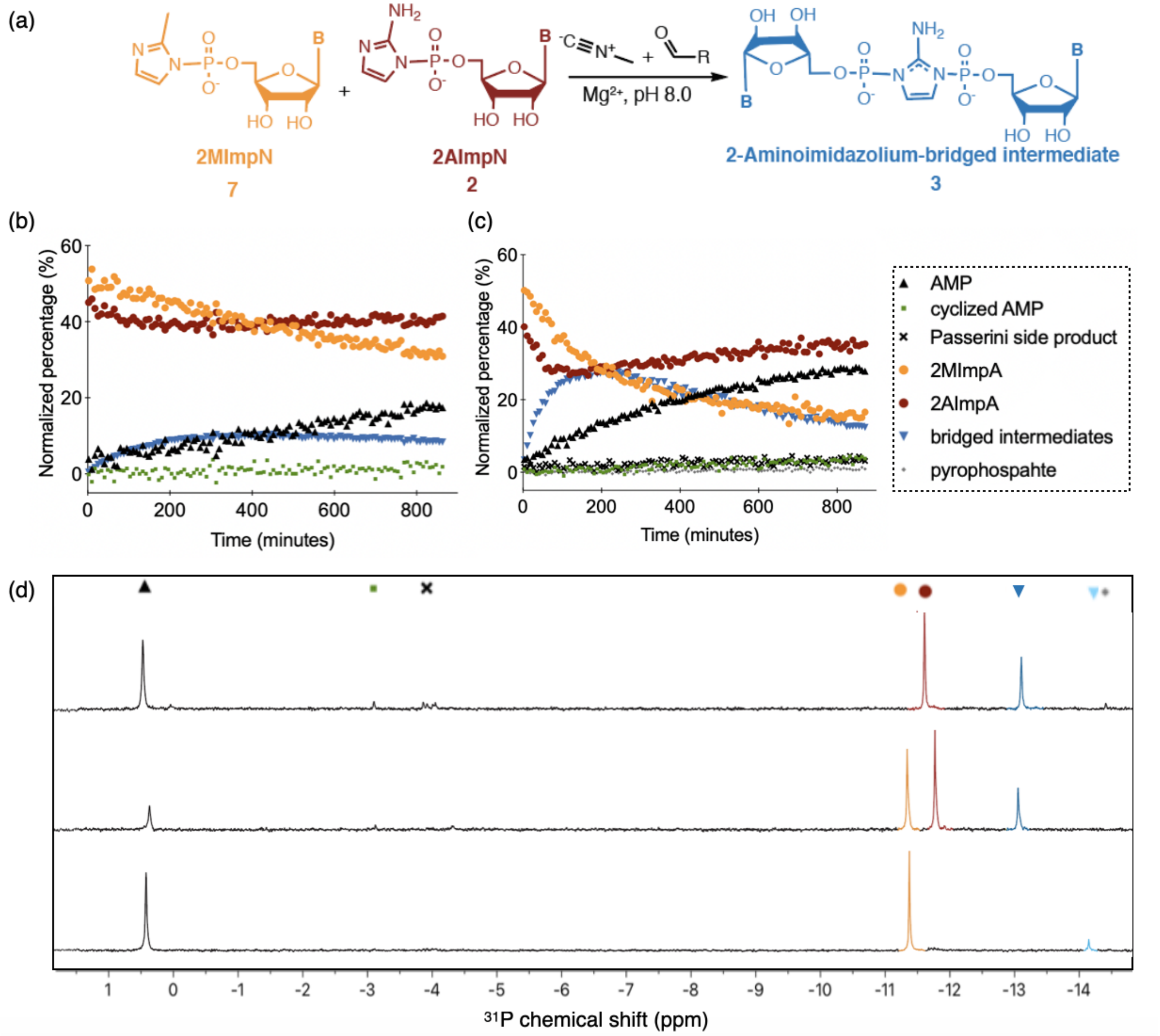

**Figure S13.** Raw PAGE data of primer extension products used for Fig. 3 under various conditions at t = 24 hours. Positions of primer and +1 to +3 products are indicated.
Reaction conditions: the corresponding amount of 2AImpNs and NMPs (as indicated below) were added to primer (1 μM, green), template (1.5 μM, black), HEPES (200 mM) pH 8, and $Mg^{2+}$ (30 mM, $MgCl_2$) with isocyanide chemistry (light blue and pink) or without (dark blue, grey, and black). n = 3 replicates.

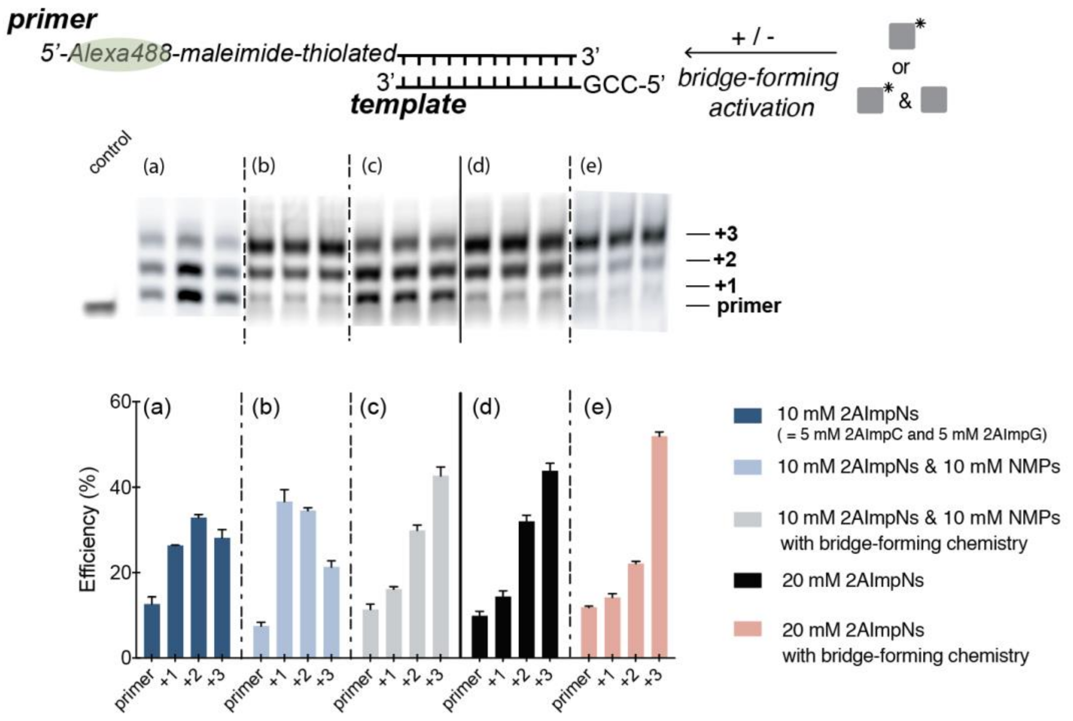

### 3. Supplementary Scheme

**Scheme S1.** Proposed mechanism for Passerini side product **5** formation.

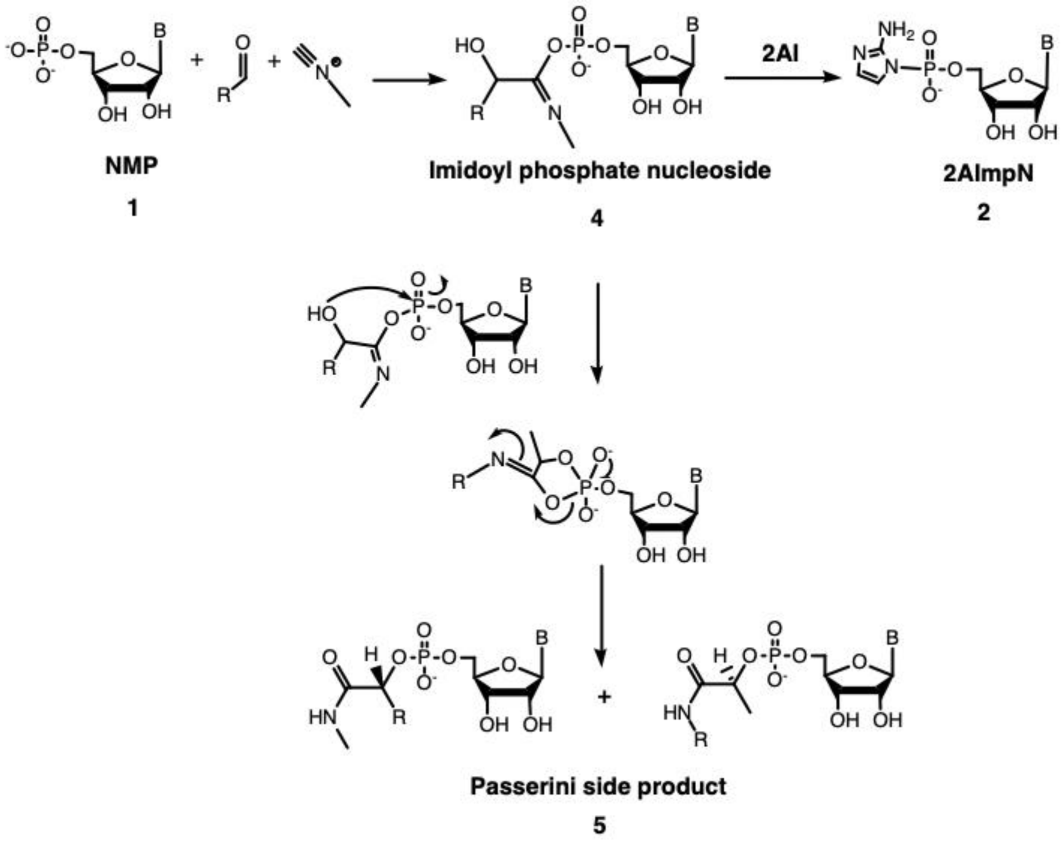

### 4. Supplementary Tables

**Table S1.** Yields and ratios of 2AImpA **2** to Passerini side product **5** across different conditions, as measured by $^{31}$P-NMR spectroscopy.

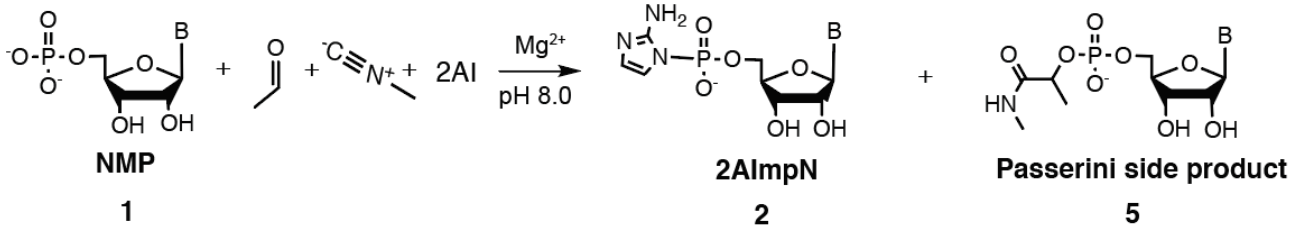


| pH | [Acetaldehyde] mM | [Methyl isocyanide] mM | [$Mg^{2+}$] mM | Buffer | [2AImpN **2**]/[Passerini product **5**] | Activation (**2**) yield % |
|---|---|---|---|---|---|---|
| | | | $Mg^{2+}$ Concentration | | | |
| 8 | 200 | 200 | 0 | HEPES | 4.6±0.5 | 31±2 |
| 8 | 200 | 200 | 10 | HEPES | 2.9±0.3 | 48±2 |
| 8 | 200 | 200 | 20 | HEPES | 2.3±0.1 | 46±1 |
| 8 | 200 | 200 | 30 | HEPES | 1.9±0.1 | 33.0±0.5 |
| 8 | 200 | 200 | 50 | HEPES | 1.2±0.1 | 32±2 |
| | | | pH | | | |
| 6 | 200 | 200 | 10 | MES | 4.0±0.4 | 38±2 |
| 7 | 200 | 200 | 10 | HEPES | 4.2±0.4 | 57±2 |
| 8 | 200 | 200 | 10 | HEPES | 2.9±0.3 | 48±2 |
| | | | Methyl Isocyanide Concentration | | | |
| 8 | 200 | 200 | 30 | HEPES | 2.3±0.4 | 36±2 |
| 8 | 200 | 300 | 30 | HEPES | 2.9±0.2 | 45±4 |
| 8 | 200 | 400 | 30 | HEPES | 2.7±0.8 | 50±4 |
| 8 | 200 | 800 | 30 | HEPES | 5.0±0.9 | 58±4 |
| | | | Acetaldehyde Concentration | | | |
| 8 | 300 | 200 | 30 | HEPES | 2.3±0.4 | 45±2 |
| 8 | 400 | 200 | 30 | HEPES | 2.9±0.6 | 50±3 |
| 8 | 800 | 200 | 30 | HEPES | 2.2±0.3 | 36±1 |
| | | | Acetaldehyde and Methyl Isocyanide Concentrations | | | |
| 8 | 30 | 30 | 30 | HEPES | 6±6 | 0.6±0.6 |
| 8 | 50 | 50 | 30 | HEPES | 5.4±0.7 | 11.9±0.1 |
| 8 | 400 | 400 | 30 | HEPES | 3.5±0.2 | 62±2 |
| 8 | 600 | 600 | 30 | HEPES | 3.7±0.3 | 71±1 |
| 8 | 800 | 800 | 30 | HEPES | 3.5±0.8 | 69±4 |

(1) All reactions were carried out using AMP (10 mM), 2AI (100 mM), and buffer (200 mM) in addition to the materials listed in the table. Errors are standard deviations of the mean, n = 3 replicates.

(2) 200 mM buffer solutions were used to maintain nearly constant pH values over a given reaction course. We note that HEPES proved an ideal buffer because it exhibited no discernible chemical reactions with the species in the activation chemistry (Table S1). The carboxylate groups of BICINE will undergo typical Passerini-type chemistry, resulting in a lower concentration of the other components of the reaction, and hence less efficient activation. Similarly, the TRIS buffer interacts with aldehyde as described previously [6].

**Table S2.** Yields and ratios of 2AImpA **2** to Passerini side product **5** with various aldehydes/ketones measured by NMR spectroscopy.

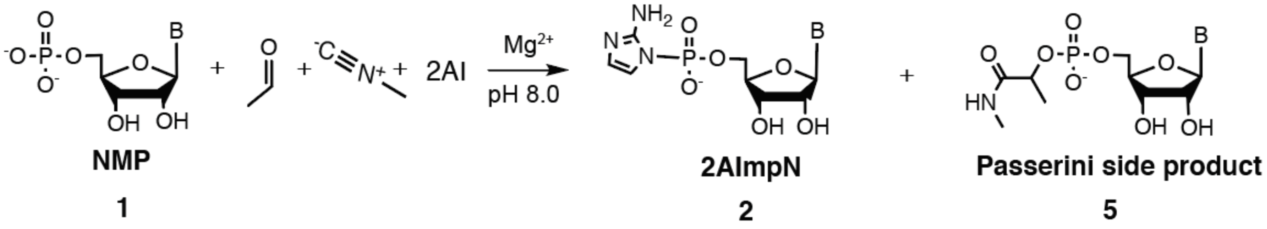


| Aldehyde or Ketone | Activation (**2**) yield% | [2AImpN **2**]/ [Passerini product **5**] |
|---|---|---|
| Glycolaldehyde | 23±1 | 2.1±0.2 |
| Acetaldehyde | 31.0±0.7 | 3.0±0.5 |
| Propionaldehyde | 50±1 | 6±1 |
| Isobutyraldehyde | 63.0±0.6 | 12.7±0.3 |
| 2-Methylbutyraldehyde | 64±2 | 18±3 |
| 2-Methylbutyraldehyde* | 89±1 | 23±3 |
| Acetone | 0 | NA |

All reactions were carried out using AMP (10 mM), aldehyde/ketone (200 mM), methyl isocyanide (200 mM), 2AI (200 mM), Mg2+ ($MgCl_2$, 10 mM), and BICINE (200 mM) at pH 8.0. Errors are standard deviations of the mean, n = 3 replicates.
*Other than BICINE being replaced by HEPES buffer, the reaction conditions were the same. A corresponding NMR spectrum for 2-methylbutyraldehyde can be found in Fig. S5.

**Table S3.** Yields and ratios of 2AImpN to Passerini side product across different conditions measured by NMR spectroscopy. All reactions were carried out using NMP (10 mM), acetaldehyde (200 mM), methyl isocyanide (200 mM), 2AI (100 mM), and buffer (200 mM) in addition to the materials listed in the table. n = 3 replicates.

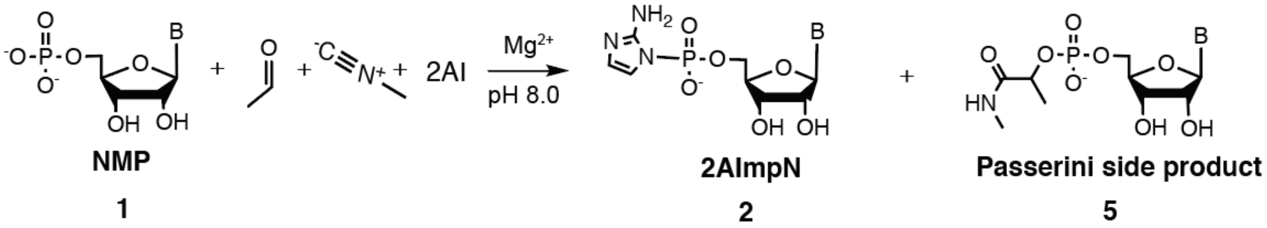


| NMP | [$Mg^{2+}$] mM | Buffer | [2AImpN]/ [Passerini product] | Activation yield % |
|---|---|---|---|---|
| | | Nucleotide Identity | | |
| GMP | 30 | HEPES | 2.2±0.3 | 39±3 |
| CMP | 30 | HEPES | 2.0±0.2 | 40±3 |
| UMP | 30 | HEPES | 1.5±0.3 | 30±3 |
| AMP | 30 | HEPES | 2.3±0.4 | 36±2 |
| | | Buffer Identity | | |
| AMP | 10 | HEPES | 2.9±0.3 | 48±2 |
| AMP | 10 | TRIS | 6±1 | 33±4 |
| AMP | 10 | BICINE | 3.0±0.5 | 31.0±0.7 |

200mM buffer solutions were used to maintain nearly constant pH values over a given reaction course. We note that HEPES proved an ideal buffer because it exhibited no discernible chemical reactions with the species in the activation chemistry (Table S1). The carboxylate groups of BICINE will undergo typical Passerini-type chemistry, resulting in a lower concentration of the other components of the reaction, and hence less efficient activation. Similarly, the TRIS buffer interacts with aldehyde as described previously [6].

**Table S4.** Sequences of oligonucleotides used in this study.

| Name | RNA Sequence (5' → 3') |
|---|---|
| control primer | /Alexa488/ AGU GAG UAA CUC |
| thiolated primer | /5ThioMC6-D/ AGU GAG UAA CUC |
| template | CCG GAG UUA CUC ACU |
| complementary | AGU GAG UAA CUC CGG |

### 5. Supplementary Text

1. Identification of the Dead-end Nucleotide Side Product
2. Development of a New Primer Extension Assay
3. Characterization of Bridge-forming Activation

#### 1. Identification of the Dead-end Nucleotide Side Product

A solution of 200 mM isocyanide, 200 mM acetaldehyde, 100 mM 2AI, and 10 mM AMP was monitored in 10% $D_2O$ by $^{31}P$ NMR over 12 hours. Besides the monophosphate starting material **1** and activated mononucleotides **2**, which were confirmed by the spike-in of their corresponding authentic standards, a pronounced nucleotide-containing side product was found (Fig. S3, S4). Isocyanides, aldehydes, and phosphates are known to undergo Passerini-type addition reactions. [7] Direct inject electrospray mass spectrometry (ESI-MS) of these reaction mixtures showed an *m/z* consistent with a Passerini product **5** (Scheme S1). Specifically, an otherwise unattributed *m/z* peak at 431.1 is found for the reaction using AMP, which is consistent with the full Passerini reaction product **5** (exact expected *m/z* = 431.11) in which intramolecular attack of the hydroxy group at the phosphorus atom is followed by a rearrangement (Scheme S1). In addition, the unique chiral center of **5** is in agreement with the observed diastereomers in the $^{31}P$ NMR spectrum (Fig. S3). The resultant amidoyl moiety of **5** is not a likely substrate for nucleophilic attack given the lack of resonance from oxygen. This is in contrast with the transient imidoyl phosphate intermediate **4** (Scheme S1). We therefore ascribe the diastereomers in $^{31}P$ NMR spectrum to the Passerini reaction product **5**, which we deem undesirable because it is not expected to contribute to RNA polymerization and may even inhibit the reaction by competing for binding sites on the template.

#### 2. Development of a New Primer Extension Assay

Typical experiments utilize a fluorescent dye-labeled primer which is then characterized by polyacrylamide gel electrophoresis (PAGE) in which different length primer extension products yield a characteristic banding pattern on the gel. However, many dyes contain functional groups such as carboxyls and amines that are expected to be incompatible with the isocyanide activation chemistry. We screened a variety of labeled primers representing multiple dye families against the activation chemistry and its constituents. In all cases, an additional band or band shift was observed due to chemical modification of the dye, in many cases probably via Passerini-type reactions (Fig. S6). We therefore sought a post-labeling strategy in which a dye is introduced after the reactions being probed have taken place. A thiol-modified primer was anticipated to be unreactive to isocyanide activation chemistry, thereby allowing the subsequent conjugation of a fluorophore-maleimide through the sulfhydryl-maleimide coupling. With this strategy, we observed no additional bands or band shifts compared to the control primer among samples that had been exposed to various combinations of isocyanide activation chemistry constituents for 24 hours (Fig. S7). Although absolute band intensities vary due to differences in purification and labeling efficiencies, only relative intensities within each lane are used to quantify the primer extension results. We next tested the accuracy of this post-labeling method by preparing a standard primer extension reaction with the template oligo 5′-CCGGAGUUACUCACU-3′, in which the template sequence 5′-CCG-3′ is copied in the presence of a primer and purified 2AImpG and 2AImpC. The quantification of primer extension using a thiol-modified primer is in very good agreement with the quantification using the standard dye-labeled primer (Fig. S8). We conclude that this post-labeling approach is an appropriate alternative technique to quantifying primer extension under otherwise dye-prohibitive conditions.

#### 3. Characterization of Bridge-forming Activation

We note that the pool of unactivated mononucleotides **1** contributes to bridged dinucleotide **3** formation because the change in activated mononucleotide **2** concentration over the course of a reaction is insufficient to account for all the formed bridged dinucleotide **3**. For example, in the case of 50:50 2AImpN:NMP (Fig. 2b), the activated mononucleotide **2** concentration decreased by 13% from t = 12 min. to t = 229 min. Therefore, a maximum of 6.5% of the bridged intermediate species **3** could come solely from the activated mononucleotides **2** (two mononucleotides being required to generate a single bridged dinucleotide **3**). (Note that the self-reaction of activated monomers **2** only formed less than 3% bridged dinucleotides **3** (Fig. 2c), which cannot account for the 6.5% increase). However, over the same time period, there was a 12% increase in the bridged dinucleotide **3** concentration. Recalling that there is no free 2AI here, these observations are consistent with bridged dinucleotides forming from (i) the activation chemistry generating an imidoyl phosphate **4** from an initially unactivated mononucleotide **1**, (ii) displacement of the imidoyl moiety by N3 of 2AI(mpN) **2**, and (iii) the background self-reaction of activated mononucleotides **2** as per the established pathway. (i) and (ii) are analogous to the isocyanide-mediated activation of mononucleotides in the presence of free 2AI, except in this case the 2AI is already attached to another mononucleotide (Fig. 1).

### 6. Supplementary References

(1) Walton, T.; Szostak, J. W., A Highly Reactive Imidazolium-Bridged Dinucleotide Intermediate in Nonenzymatic RNA Primer Extension. *J Am Chem Soc* **2016,** *138* (36), 11996-12002.
(2) R. E. Schuster, J. E. S., and Joseph Casanova, Jr., *Organic Syntheses* **1966,** *46*, 75.
(3) Li, L.; Prywes, N.; Tam, C. P.; O'Flaherty, D. K.; Lelyveld, V. S.; Izgu, E. C.; Pal, A.; Szostak, J. W., Enhanced Nonenzymatic RNA Copying with 2-Aminoimidazole Activated Nucleotides. *J Am Chem Soc* **2017,** *139* (5), 1810-1813.
(4) Mariani, A.; Russell, D. A.; Javelle, T.; Sutherland, J. D., A Light-Releasable Potentially Prebiotic Nucleotide Activating Agent. *J Am Chem Soc* **2018,** *140* (28), 8657-8661.
(5) Walton, T.; Szostak, J. W., A Kinetic Model of Nonenzymatic RNA Polymerization by Cytidine-5 '-phosphoro-2-aminoimidazolide. *Biochemistry-Us* **2017,** *56* (43), 5739-5747.
(6) Ogilvie, J. W.; Whitaker, S. C., Effect of Tris on Reactions Catalyzed by Homoserine Dehydrogenase and Glyceraldehyde-3-Phosphate Dehydrogenase. *Biochim Biophys Acta* **1976,** *445* (3), 525-536.
(7) Sutherland, J. D.; Mullen, L. B.; Buchet, F. F., Potentially prebiotic Passerini-type reactions of phosphates. *Synlett* **2008,** (14), 2161-2163.
